# CyuR is a Dual Regulator for L-Cysteine Dependent Antimicrobial Resistance in *Escherichia coli*

**DOI:** 10.1101/2023.05.16.541025

**Authors:** Irina A. Rodionova, Hyun Gyu Lim, Dmitry A Rodionov, Ying Hutchison, Christopher Dalldorf, Ye Gao, Jonathan Monk, Bernhard O. Palsson

## Abstract

Hydrogen sulfide (H_2_S), mainly produced from L-cysteine (Cys), renders bacteria highly resistant to oxidative stress. This mitigation of oxidative stress was suggested to be an important survival mechanism to achieve antimicrobial resistance (AMR) in many pathogenic bacteria. CyuR (known as DecR or YbaO) is a recently characterized Cys-dependent transcription regulator, responsible for the activation of the *cyuAP* operon and generation of hydrogen sulfide from Cys. Despite its potential importance, the regulatory network of CyuR remains poorly understood. In this study, we investigated the roles of the CyuR regulon in a Cys-dependent AMR mechanism in *E. coli* strains. We found: 1) Cys metabolism has a significant role in AMR and its effect is conserved in many *E. coli* strains, including clinical isolates; 2) CyuR negatively controls the expression of *mdlAB* encoding a transporter that exports antibiotics such as cefazolin and vancomycin; 3) CyuR binds to a DNA sequence motif ‘GAAwAAATTGTxGxxATTTsyCC’ in the absence of Cys, confirmed by an *in vitro* binding assay; and 4) CyuR may regulate 25 additional genes as suggested by *in silico* motif scanning and transcriptome sequencing. Collectively, our findings expanded the understanding of the biological roles of CyuR relevant to antibiotic resistance associated with Cys.

## 1. Introduction

The emergence of antimicrobial resistant (AMR) bacteria has been recognized as a serious threat to public health^1^. Bactericidal antibiotics (ABx, e.g., quinolones, β-lactams) generally increase the level of hydroxyl radicals resulting in lethal oxidative stress to cells^2^. It is known that hydroxyl radicals are produced from damaged key metabolic pathways including the TCA cycle, depletion of NADH, inactivated Fe-S clusters, and stimulation of the Fenton reaction^3^. Accordingly, in order to survive in the presence of ABx, bacteria utilize diverse resistance mechanisms that include changing the expression of genes responsible for oxidative stress mitigation and efflux of an ABx.

Recently, the production of hydrogen sulfide (H_2_S) was suggested to be one of the key mechanisms for increased tolerance against many antibiotics^4, 5^. Many bacteria can generate H_2_S via decomposition of L-cysteine (Cys) or reduction of an inorganic sulfur source, such as thiosulfate. It was shown that H_2_S decreases the amount of intracellular reactive oxygen species (ROS) by reacting with H_2_O_2_, a substrate of the Fenton reaction^6^, and stimulating the gene expression for enzymes that scavenge ROS. Therefore, the generation of H_2_S contributes to the enhancement of intrinsic antibiotic resistance^4^. Additionally, one recent study showed H_2_S can react with cystine and yield Cys hydropersulfide (CysSSH), which inactivates multiple antibiotics such as penicillin G, ampicillin, and meropenem^7^. These reports suggest that Cys-derived H_2_S plays a critical role in antibiotic resistance.

In *E. coli*, CyuR, also known as YbaO or DecR, was reported to be an important regulator for the generation of hydrogen sulfide from Cys in microaerobic conditions^8, 9^. Its physical interaction with the *cyuAP* operon consisting of *cyuA*, previously referred as *yhaN* or *yhaM*, and *cyuP* (b3110, also referred as *dlsT* or *yhaO*) was reported by two independent studies that utilized systematic evolution of ligands by exponential enrichment^8^ (SELEX) and chromatin immunoprecipitation exonuclease (ChIP-exo) sequencing^10^, respectively. *E. coli* has at least six enzymes (CyuA, CysM, CysK, MetC, DcyD, TnaA) that have cysteine desulfhydrase activity to generate H_2_S from decomposing Cys to pyruvate and ammonium^9, 11^. Among them, CyuA is suggested to be a major enzyme for H_2_S production in less-aerated conditions in *E. coli*^8, 9, 12^. On the other hand, *cyuP* was shown to encode an importer for Cys or serine^9, 12^. In *Yersinia ruckeri* the homologs of *cyuP and cyuA* are encoded in one operon - *cdsAB* - and involved in Cys uptake; additionally, the operon was shown to be needed for full virulence of *Y. ruckeri* in fish^13^. However, despite the importance of CyuR, much of its detailed regulatory information (e.g., its binding motif) and effects of Cys are not well characterized and thus remain unknown.

In this study, we report the effect of Cys and the detailed regulatory role of CyuR on antimicrobial resistance in *E. coli*. We utilized phenotype microarray plates to study the effects of Cys on a broad spectrum of antibiotics and growth inhibitors (hereafter collectively referred to as antibiotics) with *E. coli* laboratory strain W and three clinical isolates. Then, the role of CyuR in Cys metabolism was investigated by comparing resistance of the wildtype *E. coli* K-12 MG1655 and its *cyuR*-deletion mutant. The regulatory network of CyuR was reconstructed by investigating the clustering information of co-expressed genes (i.e., iModulons) obtained by an independent component analysis (ICA) of multiple *E. coli* gene expression profiles^14, 15^. Lastly, we show that CyuR regulation is widely conserved in many Enterobacterial genomes and for the first time report the binding motif of CyuR, enabled by genome-wide motif scanning, to predict additional target genes. Collectively, our results suggest that the elucidated CyuR-mediated L-cysteine dependent AMR mechanism is widely conserved across species and should thus be considered valuable in the treatment of AMR bacteria and Cys supplementation.

## 2. Results and Discussion

### 2.1 The presence of Cys increases antimicrobial resistance in laboratory and clinical *E. coli* strains

We investigated the effect of Cys on antimicrobial resistance in *E. coli* by using phenotype microarray (PM) plates. As a test case, we chose *E. coli* W and three clinical isolates, GN02094, GN02172, and GN02007, from the Duke Bloodstream Infection Biorepository^16^; it was confirmed that these strains generate H_2_S when Cys was supplemented at 5 mM (**Figure S1**). We grew these strains in PM11C and PM12B plates, containing 48 antibiotics at four different concentrations (**Methods**) and monitored a respiration signal, as the mean of viability of cells, in the presence and absence of 5 mM Cys (**Figure 1** and **Table 1**). Although there were variations depending on strains, notably, Cys supplementation increased respiration signals of all of these strains for 21 antibiotics (44% of the tested 48 antibiotics): L-aspartic-β-hydroxamate, chloramphenicol, cloxacillin, colistin, demeclocycline, enoxacin, erythromycin, lincomycin, lomefloxacin, minocycline, nafcillin, ofloxacin, penimepicyclin, potassium tellurite, D,L-serine hydroxamate, spectinomycin, spiramycin, sulfadiazine, sulfamethazine, tobramycin, vancomycin. Except for three antibiotics, 5-fluoroorotic acid, polymyxin B, sisomicin, increased respiration signals were observed when Cys was supplemented for the remaining antibiotics in at least one strain. These observations clearly indicate enhanced antimicrobial resistance by Cys supplementation.

**Figure 1.**
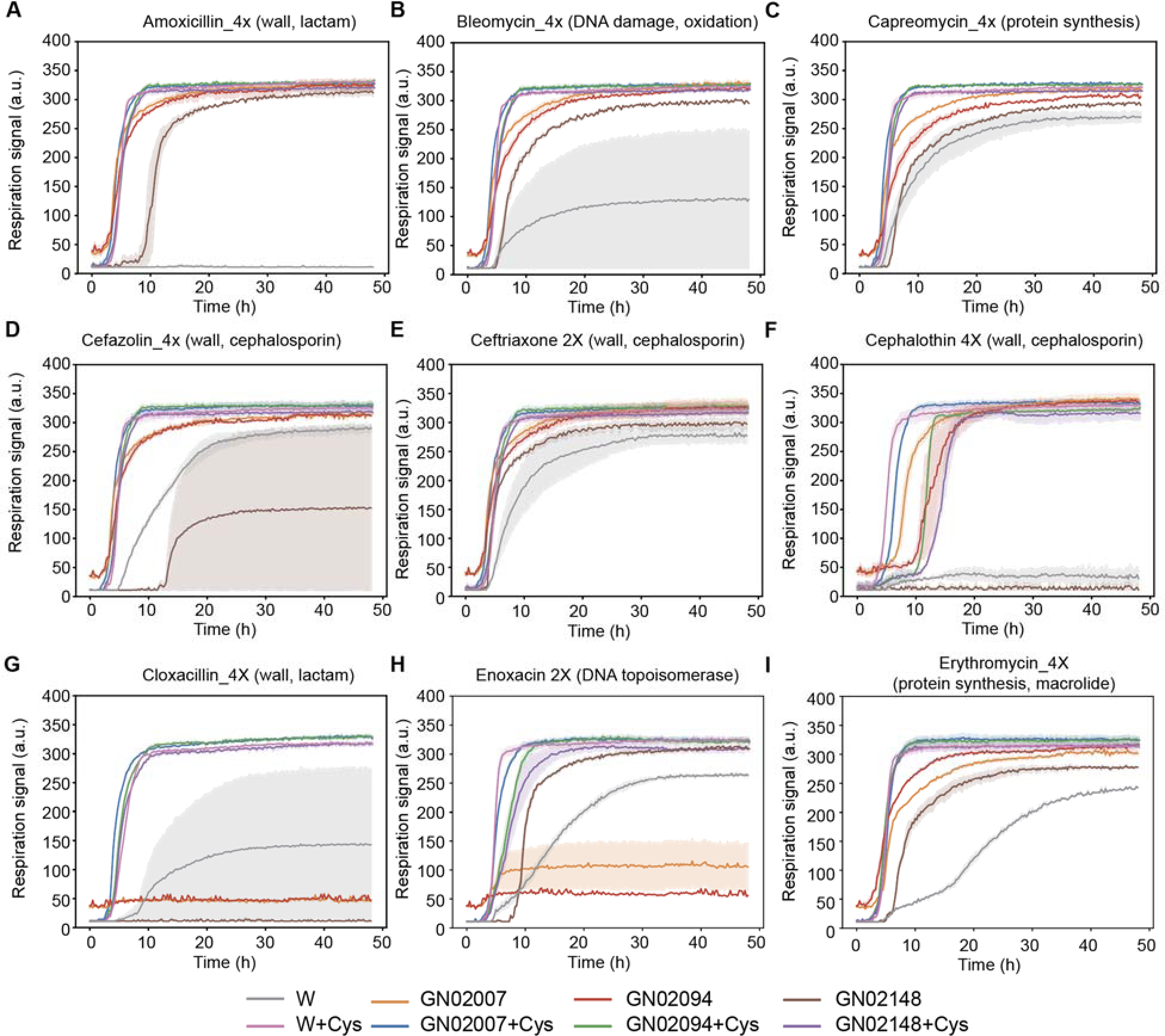
Cys supplementation increases antimicrobial resistance in *E. coli*. (**A-I**) Respiration signal of *E. coli* wild type W and three clinical isolates, GN02007, GN02094, and GN02148 in the absence and presence of 5 mM Cys with diverse antibiotics contained in phenotype microarray plates. (**A**) 4X amoxicillin, (**B**) 4X bleomycin, (**C**) 4X capreomycin, (**D**) 4X cefazolin, (**E**) 2X ceftriaxone, (**F**) 4X cephalothin, (**G**) 4X cloxacillin, (**H**) 2X enoxacin, (**I**) 4X erythromycin. In all conditions, respiration signals were high when Cys was supplemented. Results for the full antibiotics are summarized in **Table 1**.

**Table 1.**
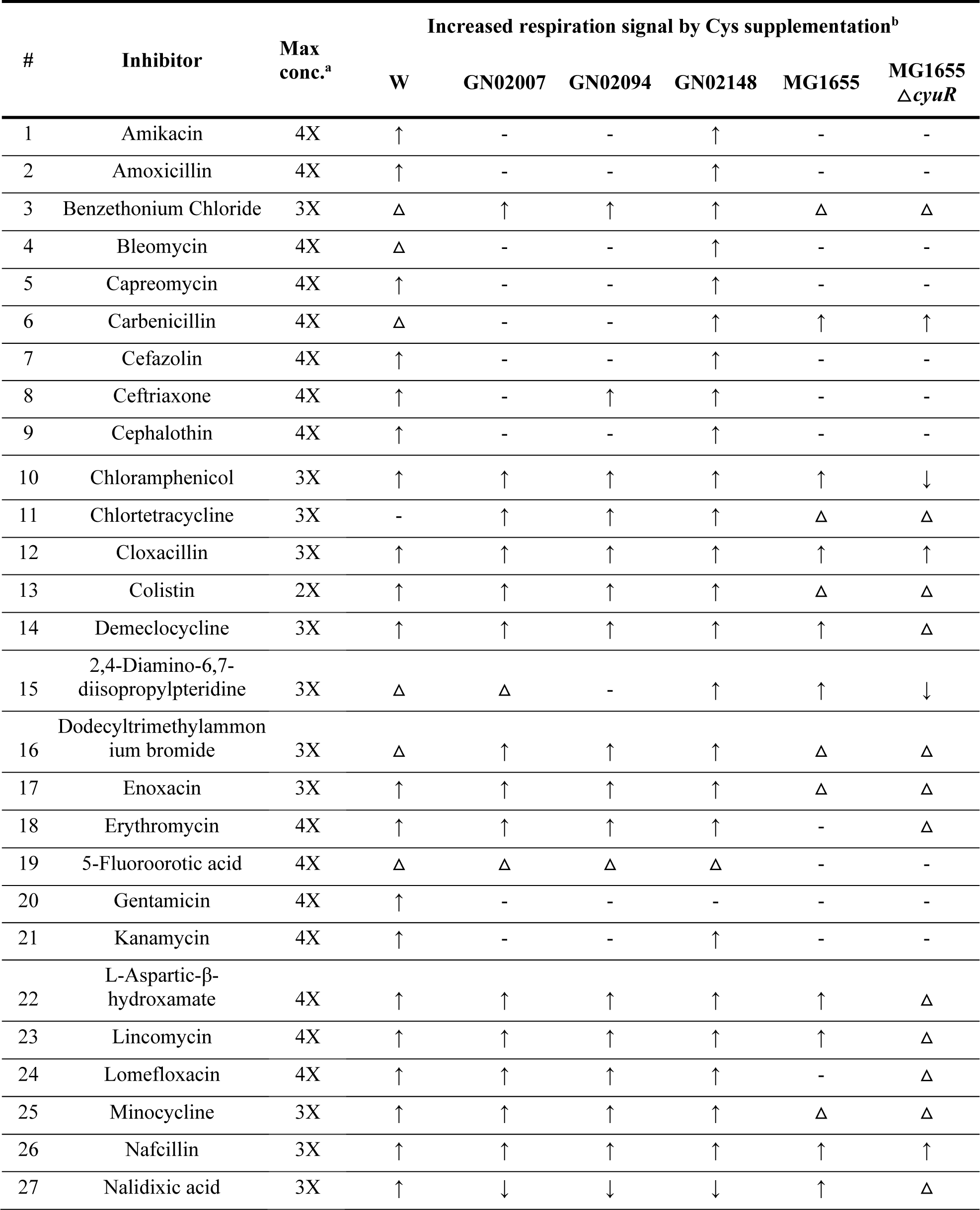
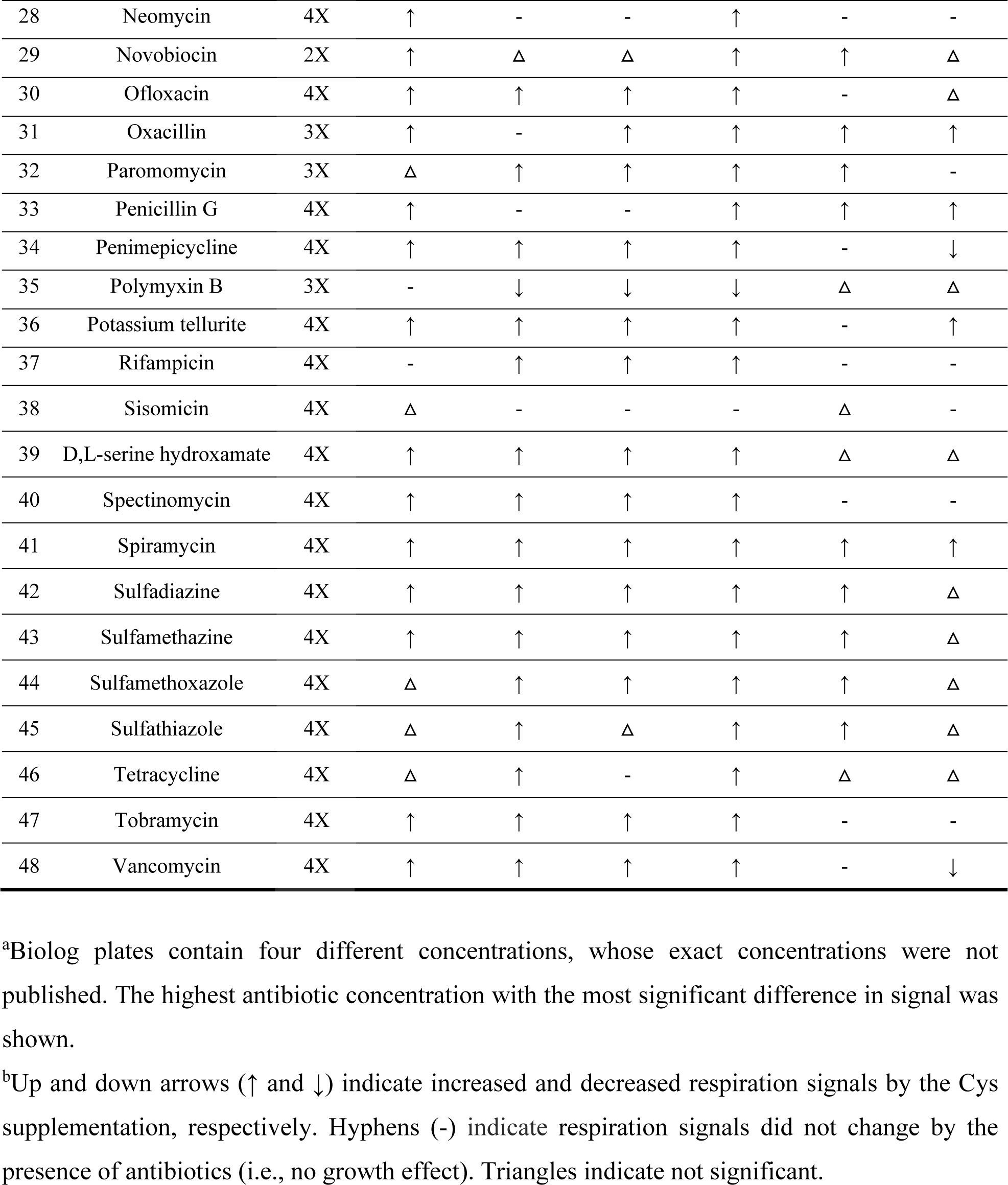
Effects of Cys supplementation to diverse growth inhibitors.

### 2.2 Transcriptomic analysis unveiled that Cys supplementation decreased stress responses in *E. coli*

To gain understanding about the increased resistance, we investigated differences in the transcriptome with Cys supplementation in *E. coli* W treated by ampicillin by analyzing iModulon activity changes (**Figure 2**). Although the conventional differential expression gene (DEG) analysis is useful, we found 2,338 genes (54.4%) out of the 4,296 annotated genes in *E. coli* with expression fold changes greater than 2 (*p*_adj_ < 0.05, **Figure S2**), challenging its comprehensive understanding. Instead, iModulons are groups of co-expressed genes, identified by performing an ICA, a machine-learning algorithm for a compendium of gene expression profiles of a given microorganism^14, 17^. It was shown that comparing iModulon activities for a dataset is effective to interpret complex transcriptome changes rather than analyzing the individual gene expression levels with a reduced dimensionality^15, 18^. For *E. coli*, a recent study identified 201 iModulons from 1,035 high quality gene expression profiles, collected under 533 conditions^14, 17^. Indeed, with this iModulon structure, a nearly half of the total variance (40.9%) in gene expression was explained and we observed that 92 iModulons had changed activities greater than 5by the Cys supplementation; the activities of 45 and 47 iModulons decreased and increased, respectively.

**Figure 2.**
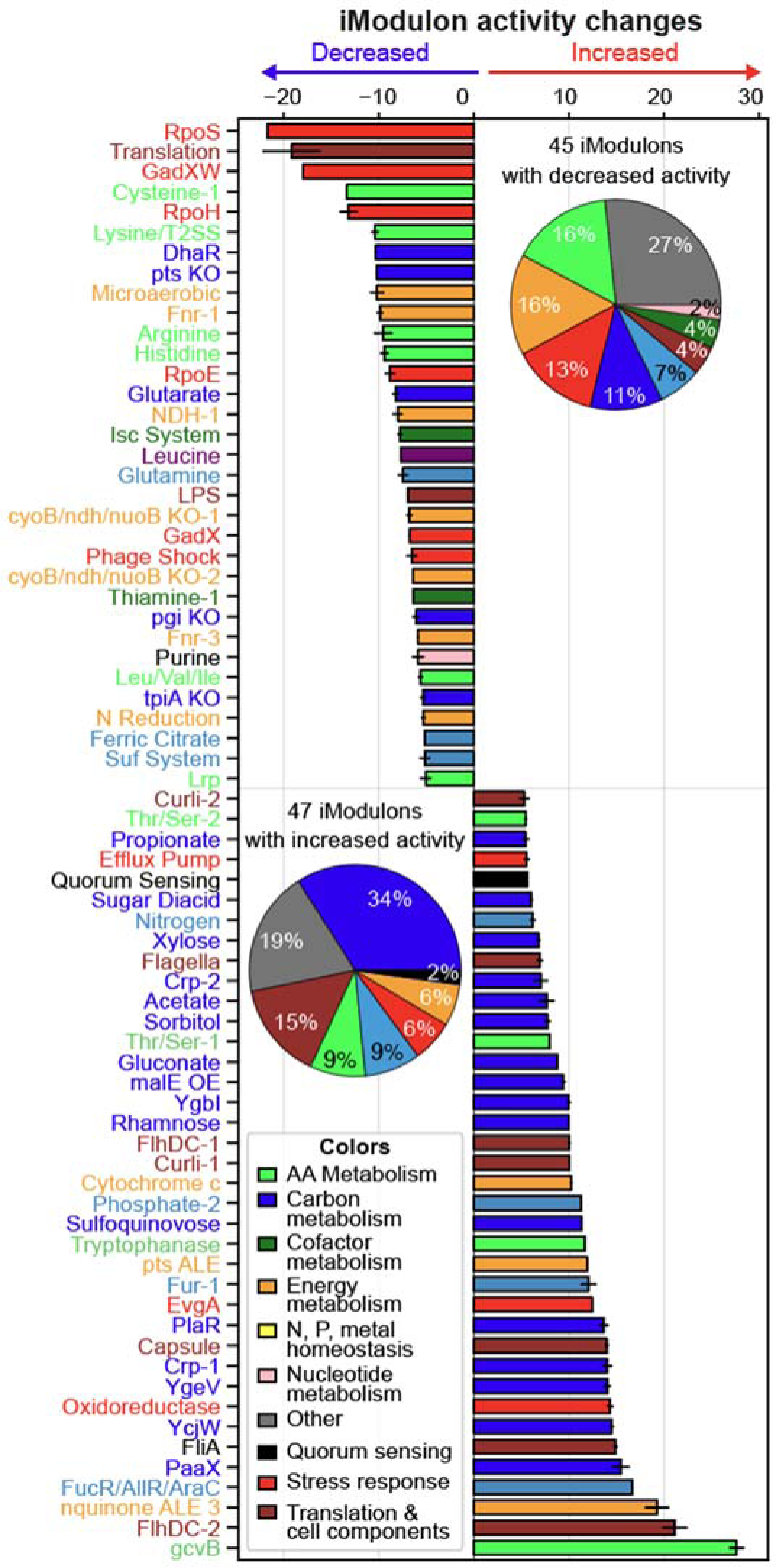
iModulon activity changes by Cys supplementation. Inferred iModulon activity changes by 5 mM Cys supplementation. The names of stress-related iModulons were colored in red. The two inset pie charts show the number of functions for iModulons with changed activities. ‘Other’ iModulons were not shown in the bar graph. A full list of iModulons and activities are in **Supplementary Table 2**. Bar colors: light green, amino acid (AA) metabolism; blue, carbon metabolism; green, cofactor metabolism; orange, Energy metabolism; yellow, nitrogen, phosphate, metal homeostasis; pink, nucleotide metabolism; grey, other; black, quorum sensing; red, stress response; brown, translation & cell components

From the changed iModulon activities, the effect of Cys supplementation was able to be inferred. Notably, many of iModulons with decreased activities include stress-response related iModulons (e.g., “RpoS” for global stress, “GadXW” for acid stress, and “RpoH” for thermal stress). This observation showed that the Cys supplementation was greatly effective to mitigate diverse stress imposed by antibiotics therefore to increase microbial tolerance. Additionally, decreased activities were observed for amino acid metabolism related iModulons (e.g., “Cysteine-1”, “Arginine”, “Histidine” for their biosynthesis), energy metabolism related iModulons (e.g., “microaerobic”, “Fnr-1”). On the other hand, many iModulons with the increased activities were related to extracellular components (e.g., “FlhDC-1”, “FlhDC-2”, and “FliA-1” for flagella, “Curli-1” for curli, “RcsAB” for colanic acid capsule, “EvgA” for the efflux pumps regulation) and carbon metabolism (e.g., “PlaR” for diketo-L-gulonate, “PaaX” for phenylacetate^19^, “YgeV” for uric acid^20^, “YcjW” for carbohydrate utilization^21^). Although detailed regulatory mechanisms are not clear, it was inferred that Cys supplementation promotes gene expression related to cellular motility and carbon catabolism, while reducing expression of stress related iModulons.

### 2.3. Inactivation of CyuR sensitizes *E. coli* to diverse antibiotics

We investigated the role of CyuR, which is known to be the major regulator for H_2_S production from Cys, to antibiotic resistance. The expression of its known target gene, *cyuA*, was highest among the six cysteine desulfhydrase genes when Cys was supplemented in the medium (**Figure S3**). We compared respiration signals of MG1655 Δ*cyuR*^10^ (**Table S1)** and its parental strain, MG1655, with and without 5 mM Cys supplementation in PM11C and PM12B plates (**Figure 3**). Similar to *E. coli* W strain, the MG1655 strain showed increased respiratory signals when Cys is present for 17 of the 48 tested antibiotics (35%). Although Cys supplementation was effective to the less number of antibiotics than those effective for *E. coli* W, this observation strongly supports similar stress reduction by the generation of H_2_S in *E. coli* MG1655.

**Figure 3.**
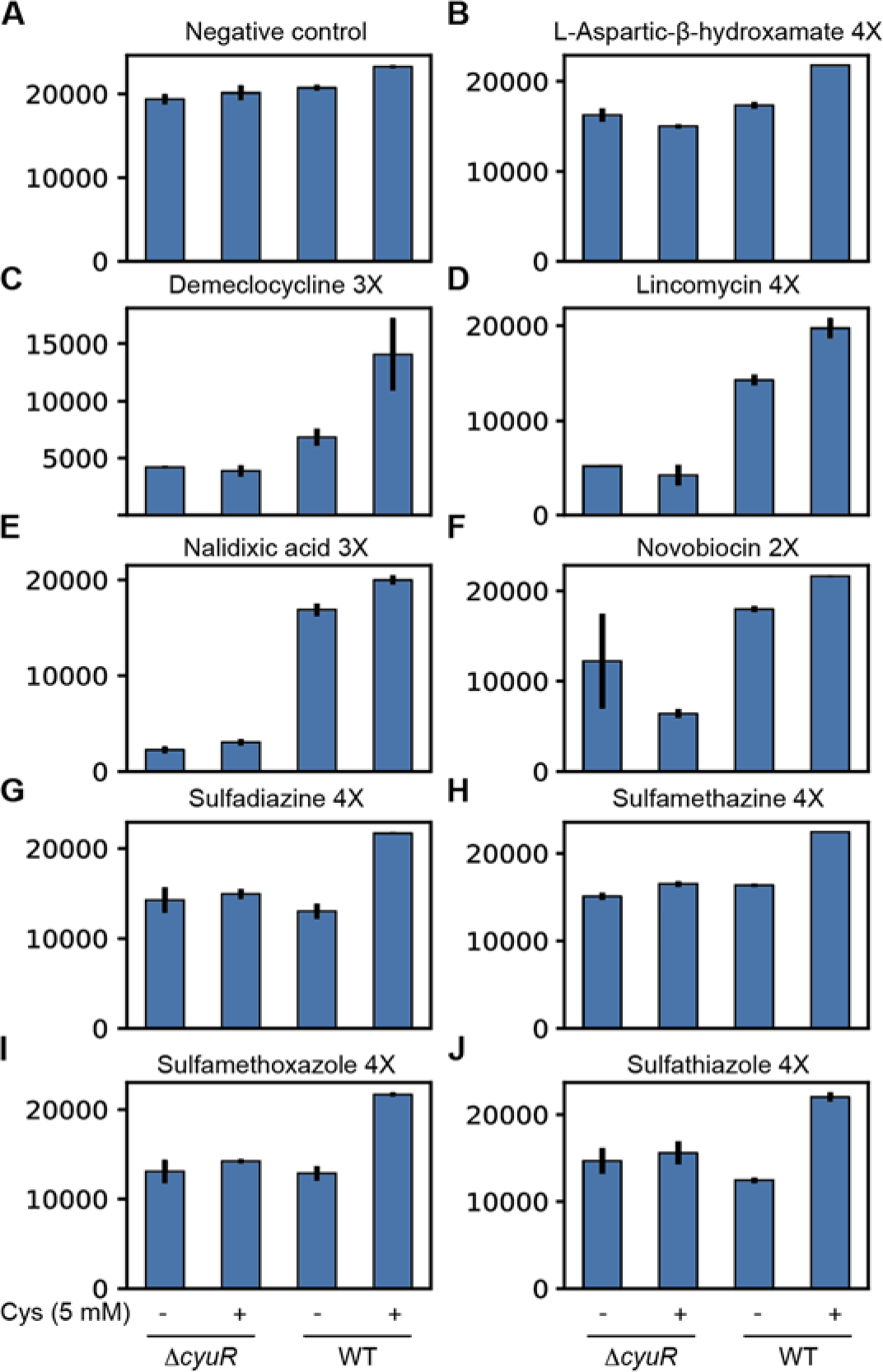
The essentiality of CyuR for Cys-involved antimicrobial resistance. (**A-J**) Respiratory signal (arbitrary unit, a.u.) of *E. coli* K-12 MG1655 and its *cyuR*-deleted mutant grown in a Biolog plate with nine different antibiotics: (**A**) L-aspartic-β-hydroxamate 1X (negative control), (**B**) L-aspartic-β-hydroxamate 4X, (**C**) demeclocycline 3X, (**D**) lincomycin 4X, (**E**) nalidixic acid 3X, (**F**) novobiocin 2X, (**G**) sulfadiazine 4X, (**H**) sulfamethazine 4X, (**I**) sulfamethoxazole 4X, (**J**) sulfathiazole 4X. *y*-axis indicate respiration signal (a.u.).

Notably, such improvements by Cys supplementation were abolished after deleting *cyuR* in the MG1655 strain for at least nine antibiotics (**Figure 3A-J**), implying the essentiality of CyuR in Cys-involved antimicrobial resistance. These antibiotics are demeclocycline, L-aspartic-β-hydroxamate, lincomycin, nalidixic acid, novobiocin, paromomycin, sulfadiazine, sulfamethazine, sulfamethoxazole, and sulfathiazole. Interestingly, the resistance was decreased by the deletion of *cyuR* itself, even when Cys was not supplemented in some cases (e.g., lincomycin, nalidixic acid). This observation suggested that CyuR is potentially involved in an additional ABx resistance mechanism, not related to Cys.

### 2.4 CyuR is a dual regulator for *cyuR*-*mdlAB* and *cyuAP* in response to Cys exposure

Given its potential expanded regulatory roles in antimicrobial resistance, we further carried out a detailed characterization of CyuR. We first gathered existing information about the expression of *cyuR* in *E. coli* and verified it using an *E. coli* gene expression compendium^15^. Previous studies^22, 23^ reported binding sites of SoxS and MarA, responsible for oxidative stress mitigation^22^ and antibiotic resistance^24^ in the upstream region of the *cyuR* gene (**Figure 4A**), implying its relevance to the stress resistance. Indeed, the expression of *cyuR* is highly correlated to the expression of 117 genes, mostly regulated by SoxS (the SoxS iModulon^15, 25^, **Figure 4B**). The expression level of *soxS* and *cyuR* was also highly correlated (pearson R of 0.57, **Figure S4A**), although the correlation between *cyuR* and *marA* was not high (pearson R of 0.16, **Figure S4B**). These observations confirmed that CyuR is under regulation by SoxS and that it is an important regulator for mitigating various types of stress.

**Figure 4.**
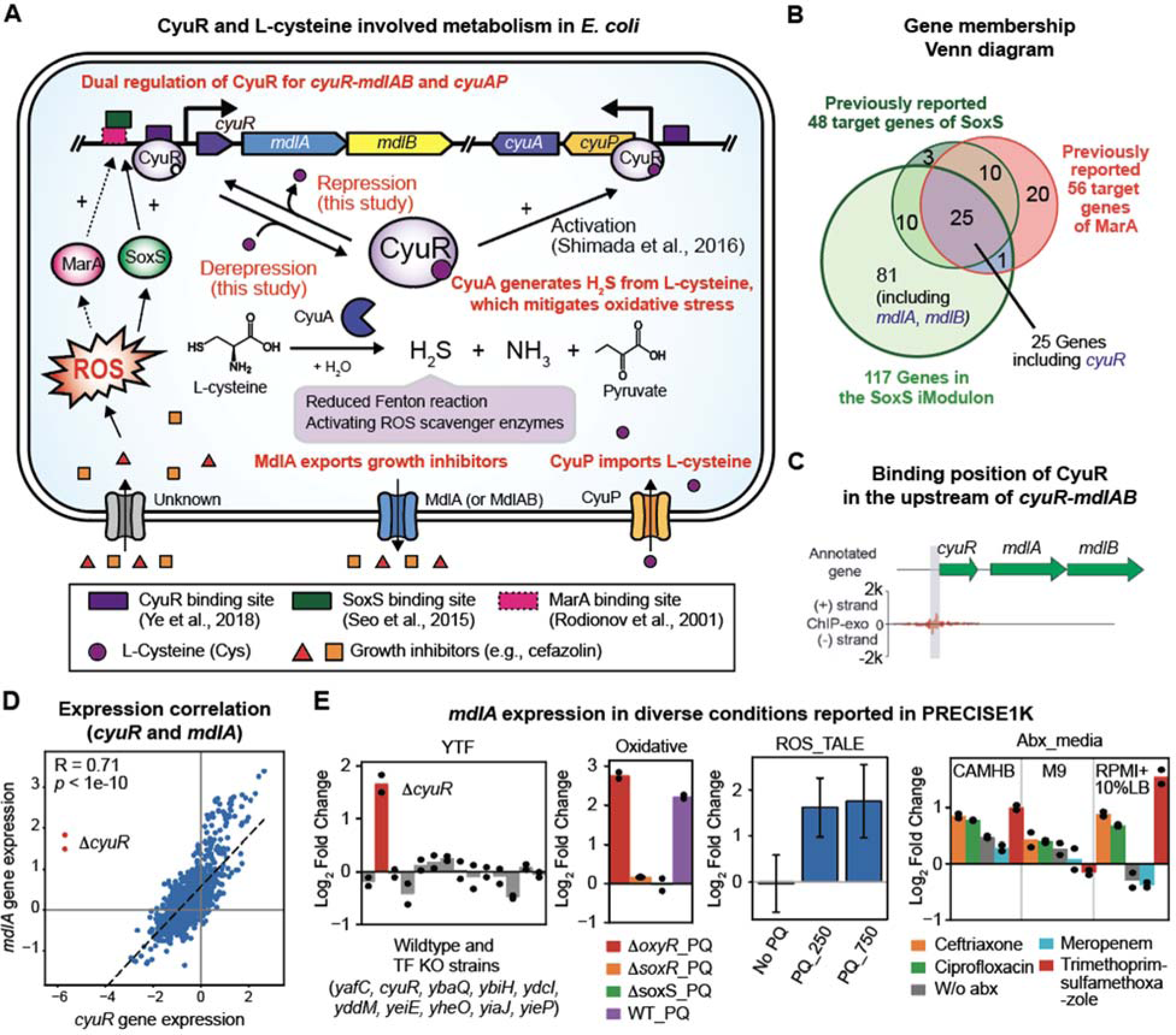
Regulatory information of CyuR and its regulon genes. (**A**) CyuR (b0447, also called YbaO, DecR) and L-cysteine (Cys) involved metabolism in *E. coli*. CyuR is a dual regulator for *cyuAP*^8^, encoding Cys sulfifdrase and importer, and *mdlAB*, encoding a novel antibiotic efflux ABC-type transporter MdlA (or MdlAB). In the presence of Cys, Cys-bound CyuR activates the expression of *cyuAP* and derepresses the expression of *mdlAB*. Binding sites of CyuR were obtained from previous studies^8, 10^. Degradation of Cys by CyuA generates hydrogen sulfide (H_2_S) which is beneficial to the reduction in Fenton reaction or stimulation of reactive oxygen species scavenging pathway. The expression of *cyuR* is regulated by SoxS^22^ and potentially MarA^23^. (**B**) Binding site of CyuR identified from a previous Chip-exo experiment^10^. *y*-axis shows the number of reads mapped to a region. (**C**) A Venn diagram of three gene groups: SoxS iModulon, SoxS regulon, MarA regulon. *cyuR* belongs to the three gene groups; *mdlA* and *mdlB* only belong to SoxS iModulon; *cyuA* and *cyuP* were not included in any groups. (**D**) Gene expression correlation between *cyuR* and *mdlA*. Expression of these two genes is highly correlated, supporting that they are co-expressed from the same operon (**E**) Gene expression of *mdlA* in multiple projects of PRECISE-1K^15^. The titles of the bar graphs indicate the names of projects that generated gene expression profiles. YTF, a project characterized the regulons of putative transcriptional factor in *E. coli*^10^; Oxidative, a project characterized the regulon of OxyR, SoxS, and SoxR^22^; ROS_TALE, a project studied a wildtype strain and evolved strains tolerized to reactive oxygen species; Abx_media: a project studied transcriptional responses to multiple antibiotics in different media conditions^42^.

We then further explored the regulation by CyuR in *E. coli* by revisiting a previously reported ChIp-Exo dataset^10^ for CyuR and the iModulon analysis of the *E. coli* gene expression compendium^15^. So far, two binding sites were reported; one binding site in the upstream of *cyuAP* was thoroughly validated whereas the other binding site in the upstream of *cyuR* itself was not validated (**Figure 4C**). Interestingly, there are two genes *mdlAB*, encoding putative transporters, co-localized with *cyuR* and they were also included in the SoxS iModulon, suggesting their co-expression. Indeed, the correlation between *cyuR* and *mdlA* was high (pearson R of 0.71 for all 1,035 samples or 0.77 for all samples except the *cyuR*-deleted strain, **Figure 4D**).

Since there was no predicted transcriptional start site in the intergenic region of *cyuR* and *mdlA*, it was inferred that they form an operon. The *mdlA* gene was differentially up-regulated when *cyuR* was deleted and the oxidative-stress-inducing paraquat (PQ) was added (the YTF, Oxidative, ROS_TALE projects, **Figure 4E**). Furthermore, the addition of several antibiotics also up-regulated this gene in the CAMHB and RPMI+10% LB medium, but not in the M9 medium (the Abx_media project, **Figure 4E**). Its up-regulation was not observed when either *soxS* or *soxR* was deleted, supporting their regulatory roles for *mdlA* (and *cyuR*). In addition, since a deletion of *cyuR* up-regulated the *mdlAB* expression, CyuR was inferred to be an autorepressor. We also further investigated the correlation between *cyuR* and *cyuP* expression (**Figure S4C**). Interestingly, the expression of the two genes did not significantly correlate (pearson R of 0.14), likely due to absence of the effector of CyuR in most samples. Cys was previously suggested to activate the DNA-binding for CyuR for the activation of *cyuPA* operon. The CyuR DNA-binding motif upstream of *cyuPA* was not predicted, but the CyuR binding was shown. Collectively, these observations confirmed that CyuR a Cys-responsive regulator for both *cyuR*-*mdlAB* and *cyuAP*.

### 2.5. Identification of the binding motif of CyuR

We determined the binding motif of CyuR by utilizing a phylogenetic footprinting assay^26^ and validated by *in vitro* fluorescence polarization assay. We hypothesized that the CyuR binding site can be predicted from the alignment of the upstream regions of c*yuR*-*mdlAB* from diverse species, given it was found that *cyuR-mdlAB* is highly conserved in many Proteobacteria (**Figure S5**); it is commonly present in *E. fergusonii*, *E. albertii, Shigella boydii, S. dysenteriae, S. sonei, Citrobacter freundii, C. youngae, C. koseri*, *C. farmeri, C. braakii, Salmonella bongori, S. enterica, Enterobacter asburiae*. When we aligned the *cyuR* upstream regions in 22 Enterobacteria, a conserved 23-bp sequence was identified, likely suggesting it is a putative binding site of the CyuR regulator (**Figure 5A**, yellow box), which was consistent with a previously reported binding site identified via Chip-exo (**Figure 4C**). The common motif is an imperfect palindrome with consensus **G**AAw**AAAT**TGTxGxx**ATTT**syC**C**, where ‘x’ is either T or C, ‘s’ is either A or C, ‘y’ is either A or G, and ‘w’ is either A or T (**Figure 5B**). The transcription of *cyuR* was predicted to be RpoD sigma factor dependent^27^.

**Figure 5.**
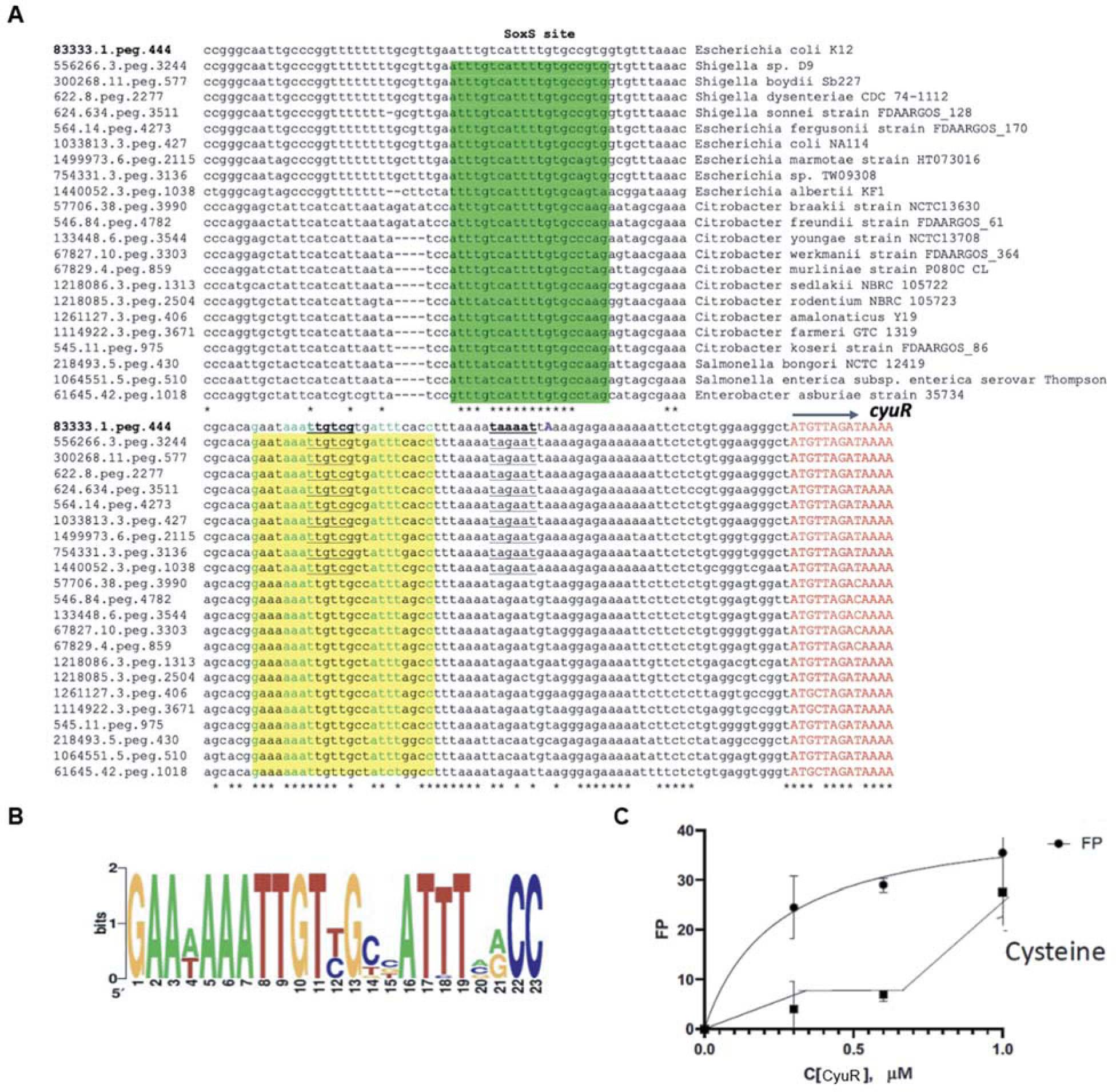
Sequence alignment of the upstream regions of CyuR in various microorganisms in the family of *Enterobacteriaceae*. (**A**) Sequence alignments of *cyuR* regions in different microorganisms. Known SoxS binding motif and predicted CyuR binding motif were colored in green and yellow, respectively. Asterisk indicates the commonality of corresponding bases. Each sequence starts with a feature id that was given by the SEED database (https://www.theseed.org). (**B**) Sequence logos were generated based on the multiple alignments of CyuR binding sites from multiple groups of *Enterobacteria* genomes (see **Figure S1**) using the WebLogo tool (https://weblogo.berkeley.edu/logo.cgi). (**C**) The fluorescence polarization of CyuR and 10 nM fluorescently labeled 30 bp DNA fragment of the upstream sequence of the *cyuR* (see **Methods**). This assay was performed in the presence and absence of 15 mM L-cysteine. *x*-axis and *y*-axis indicate CyuR concentration (μM) and fluorescence polarization, respectively.

To confirm the binding of CyuR to the predicted motif, we conducted a fluorescence polarization assay^28^ in the absence and presence of Cys (**Figure 5C**, see Methods). Purified CyuR was incubated at different concentrations (0, 0.3, 0.6, 1.0 μM) with 10 nM fluorescently labeled 30 bp DNA fragments of the upstream sequence of the *cyuR* gene. Notably, after an 1-h incubation, we observed that the FP signal increased as the amount of CyuR increased. This observation confirmed that the effector of CyuR is Cys and it indeed binds the predicted consensus sequence. The affinity of the CyuR binding to the DNA was impaired at 15 mM Cys.

### 2.6. MdlA, which is co-expressed with CyuR, is a putative efflux pump for antibiotics

We hypothesized that MdlA or MdlB can function as an antibiotic resistance efflux pump because *mdlAB* were predicted to encode membrane proteins with a nucleotide-binding domain. This domain is highly conserved in ATP-binding cassette antibiotic efflux pumps^29^. Previously, a heterologous expression of SmdA, a homologous transporter to MdlA (an identity of 79%), from *Serratia marcescens* in *E. coli*, increased resistance against norfloxacin, tetracycline, and 4,6-diamidino-2-phenylindole^30^. This hypothesis was further supported by the fact that up-regulation of *mdlA* expression was observed in the presence of trimethoprim with sulfamethoxazole, ciprofloxacin and ceftriaxone in the RPMI medium (**Figure 4E**).

We investigated their role using phenotype array plates. To test whether MdlA or MdlB can export antibiotics, we took BW25113, BW25113 Δ*mdlA*, and BW25113 Δ*mdlB* strains from the Keio collection (**Table S1**) and compared their phenotypes (**Figure 6A-F**) using 11C and 12B plates in the IF10b medium in the presence of Cys. BW25113 has the exact same sequences for *cyuR*, *mdlA*, and *mdlB* in *E. coli* K-12 MG1655. Notably, we found a difference in respiration signals from these strains in the presence of two antibiotics, cefazolin and vancomycin. Although the three strains did not show significant differences at low concentrations, the BW25113 Δ*mdlA* strain displayed a much longer lag period before growth at the 4X conditions, suggesting that MdlA is important for their efflux. Inconsistent with the previous study with *S. marcescens*^30^, no significant growth difference was observed with different levels of tetracycline (data not shown), likely because the specificity of MdlAB homologous transporters can be different. Interestingly, the deletion of *mdlB* did not affect the viability in the presence of cefazolin. Although SmdA and SmdB in *S. marcescens* were suggested to form a heterodimer transporter^30^, it is likely that MdlA and MdlB are independent to each other or MdlA can be solely active. Nevertheless, these results showed that *mdlA*, in the *cyuR-mdlAB* operon, encodes an efflux pump for antibiotics, at least for cefazolin and vancomycin.

**Figure 6.**
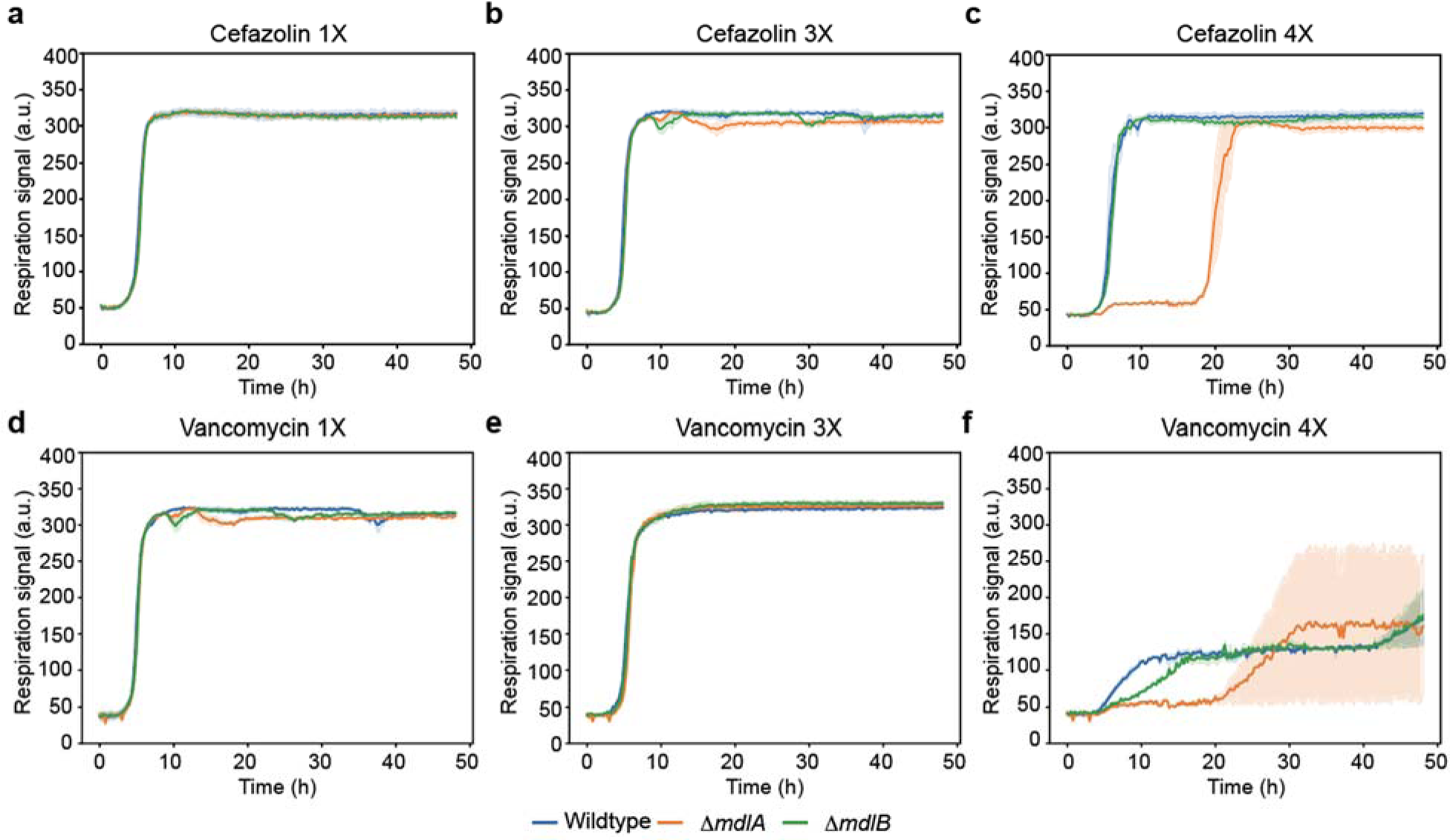
Phenotype microarray data for BW25113, BW25113 Δ*mdlA*, and BW25113 Δ*mdlB* in the presence of three different concentrations of cefazolin and vancomycin. (**A-F**) Time-course signals of BW25113 (wildtype, blue line), BW25113 Δ*mdlA* (orange line), and BW25113 Δ*mdlB* (green line) grown in a Biolog plate at the three different concentrations (1X, 3X, 4X) of (**A-C**) cefazolin and (**D-F**) vancomycin. *x*-axis and *y*-axis indicate time (h) and respiration signal (a.u.), respectively.

### 2.7. Motif scanning suggests additional target genes of CyuR

We further explored a possibility that CyuR regulates additional target genes by motif scanning of the genome of the *E. coli* MG1655 and comparative gene expression analysis (see **Methods**). From this analysis, additional putative binding sites were identified in the upstream region of 25 genes (**Table 2**). For these genes, we investigated gene expression changes by the deletion of *cyuR*^10^ or supplementation of Cys. Notably, the expression levels of four genes, *setB* encoding sugar export transporter, *aldB* encoding aldehyde dehydrogenase, *malE* and *malK* encoding maltose ABC transporter, were commonly up-regulated in the two conditions. Considering their gene expression patterns, they can be repression targets by CyuR in the absence of Cys. Additionally, there were a number of genes that were up-regulated by the supplementation of Cys. For example, *yaiV*, a DNA-binding regulator gene involved in resistance to oxidative stress^31^, *etp* encoding a lipid A modification gene, and *lapA* encoding a lipopolysaccharide assembly encoding gene were up-regulated, suggesting that these genes are potential up-regulated by CyuR when Cys is present. Taken together, these results imply that CyuR may have further expanded regulatory roles for diverse metabolism in *E. coli*.

**Table 2.**
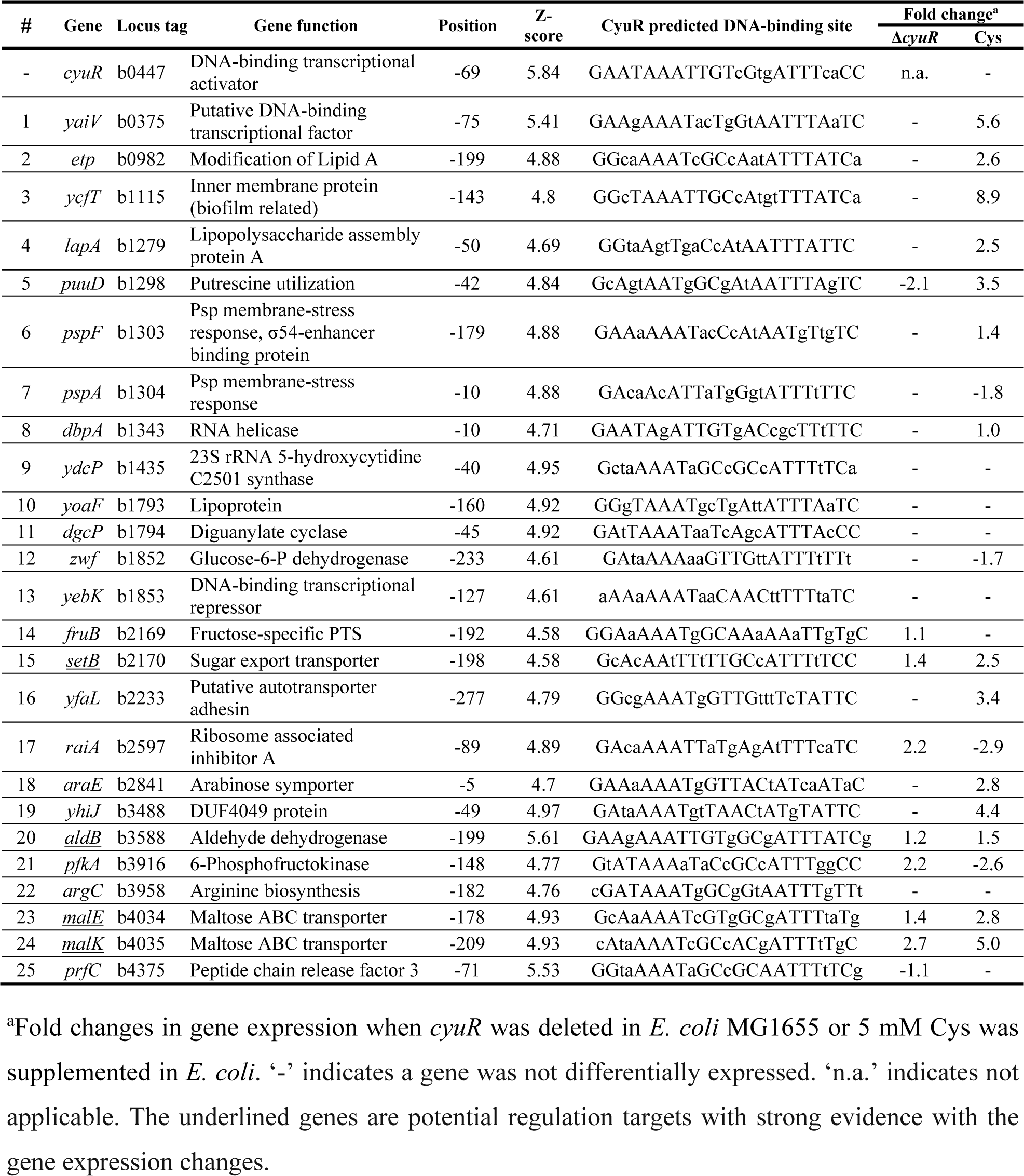
Potential regulation targets of CyuR, suggested by motif scanning.

## 3. Conclusions

In this manuscript, we showed that: 1) Cys supplementation led to the generation of H_2_S, which increases antimicrobial resistance by reducing various stress responses, supported by transcriptomic analysis using iModulons; 2) CyuR is an important regulator for Cys-dependent resistance for at least nine antibiotics: demeclocycline, L-aspartic-β-hydroxamate, lincomycin, nalidixic acid, novobiocin, sulfadiazine, sulfamethazine, sulfamethoxazole, sulfathiazole; 3) CyuR is an autorepressor for itself and antimicrobial efflux pump genes *mdlAB*, which likely forms an operon; 4) CyuR binds to a palindromic conserved motif “GAAwAAATTGTxGxxATTTsyCC”, identified by a phylogenetic footprinting assay and verified by a fluorescent polarization assay; and 5) CyuR may have more expanded regulatory roles in diverse cellular metabolism, inferred from 25 putative additional target genes, identified by motif scanning and transcriptome sequencing. Given that Cys is widely utilized as food supplement and it increased antimicrobial resistance, our study suggests that its potential harmful effect of Cys should be studied when treating bacterial infection diseases.

## 4. Materials and methods

### 4.1 Bacterial strains and growth conditions

A list of the strains used in this study was listed in **Table S1**. MG1655 *cyuR* was generated in a previous study^10^. For a routine cell culture, cells had been initially grown overnight in LB medium. The cells were refreshed for 3 hours in fresh LB media. These cultures were washed and inoculated again into a fresh M9 medium with 4 g/L glucose in the presence or absence of L-cysteine. This preculture was then diluted back to OD_600_ 0.01 into a main culture medium.

### 4.2. Phenotype microarray

*E. coli* strains were phenotyped by using microarray plates and an Omnilog system purchased from Biolog (https://www.biolog.com, Hayward, CA). These strains were inoculated from a corresponding glycerol stock to an LB medium. The next day, the cultures were re-inoculated in an IF10b medium and further incubated overnight. Then, the cultures were washed, diluted at recommended OD_600_ (approximately 0.08) in the IF10b medium, and incubated in the 11C or 12B microplates containing diverse antibiotics at four different concentrations at 37℃. Bacterial respiration was monitored by using Biolog-redox dye with or without a supplementation of 5 mM Cys with two biological replicates. It should be noted that the exact concentrations of each antibiotic are not publicly available and only relative concentrations were available.

### 4.3 Transcriptome sequencing (RNA-Seq)

*E. coli* W was grown in the presence and the absence of Cys. Pre-cultures were prepared in the LB medium. Overnight cultured cells were washed twice with the M9 medium and inoculated at an OD_600_ of 0.05 in the same medium. RNA sequencing data were generated under aerobic exponential growth conditions in M9 medium and supplemented with 5 mM Cys. To induce oxidative stress, we included 15 μg/mL of ampicillin^2^. The cells were collected at an OD_600_ of 0.6 and treated using the Qiagen RNA-protect bacterial reagent (Hilden, Germany) according to the instruction from the manufacturer. Pelleted cells were stored at −80°C, and after cell resuspension and partial lysis, they were ruptured with a beat beater. Subsequently, the total RNA was extracted using a Qiagen RNA purification kit. After total RNA extraction, ribosomal RNA was removed^32^ and the quality was assessed using an Aglient Bioanalyser using an RNA 6000 kit. Paired-end RNA sequencing libraries were prepared by using a KAPA RNA HyperPrep kit from Kapa Biosystems (Wilmington, MA, USA) as described previously^33^. Raw-sequencing reads were aligned to the reference genome of the *E. coli* MG1655 strain (accession number: NC_000913.3) using Bowtie2^34^. Transcripts per million calculation and differentially expressed gene analysis were performed by DESeq2 (v1.22.1)^35^. iModulon activity analysis was performed by using the Pymodulon package (https://github.com/SBRG/pymodulon)14.

### 4.4 Prediction of the binding site of CyuR and motif scanning

The potential CyuR-binding site was identified by a phylogenetic footprinting approach using multiple sequence alignments. Orthologs of the *E. coli cyuR* gene in other Proteobacteria, as well as multiple sequence alignment of orthologous upstream region, was identified using the PubSEED comparative genomics platform^36^. CyuR-binding site logos were obtained by using the WebLogo tool^37^. The aligned CyuR sites from *E. coli* and other proteobacteria were used as a training set to build a Positional Weight Matrix (PWM) Positional nucleotide weights in the CyuR-PWM and Z-scores of candidate sites were calculated as the sum of the respective positional nucleotide weights.

Genome scanning for identifying additional candidate CyuR-binding sites was performed using the Genome Explorer software^38^. The threshold for the site search was defined as the lowest score observed in the training set (Z-score = 4.5).

### 4.5 *In vitro* DNA binding assay for CyuR

A single colony of the *cyuR* overexpressing strain of *E. coli* obtained from the ASKA collection^39^, was inoculated into a LB medium from an agar plate containing chloramphenicol and grown at 37℃. After overnight, the culture was further passaged to a 50 mL fresh medium. After OD_600_ reached 0.8, 0.8 mM isopropyl β-D-1-thiogalactopyranoside was added. The cultures were incubated at 24 °C overnight with continuous shaking, and cells were collected by centrifugation and lysed as previously described^40^. The CyuR recombinant protein containing an N-terminal 6хHis tag was purified by Ni-chelation chromatography from the soluble fraction as described^40, 41^. The insoluble fraction was solubilized in 7 M urea At-buffer and purified on a Ni-NTA mini-column (Qiagen Inc.) with At-buffer (100 mM Tris-HCl buffer, pH 8, 0.5 mM NaCl, 5 mM imidazole, and 0.3% Brij, β-mercaptoethanol with 7 M urea. The expression of a proper size of CyuR was confirmed by performing a SDS-PAGE analysis.

The interaction of the purified CyuR protein with its specific DNA-binding site was assessed using fluorescent polarization with fluorescently labeled 28-bp orthologous DNA fragment from *Citrobacter braakii* (5’-a<u>ggAAAATTGTTGCCATTTAGCCTTTAAAATgga</u>), containing the predicted orthologous CyuR-binding site and flanked on each side by extra guanine residues. For each DNA fragment, two complementary single-stranded oligonucleotides were synthesized by IDT, at which the fragments were labeled by 6-carboxyfluorescein at 5’ end. The double-stranded DNA fragments were obtained by annealing the labeled oligonucleotides with unlabeled complementary oligonucleotides at a 1:10 ratio. The obtained fluorescently labeled DNA fragments (10 nM) were incubated for 1 hour at 30°C with the increasing concentrations of recombinant CyuR protein in the assay mixture (0.1 mL) in 96-well black plates. The binding buffer contained 20 mM Tris-HCl (pH 7.5), 0.1 M NaCl, 0.5 mM EDTA, 10 mM MgSO_4_, 2 mM DTT, 5 μg/mL herring sperm DNA. The fluorescence-labeled DNA was detected by using a Beckman Coulter DTX 880 Multimode Detector. We monitored the fluorescent polarization in the absence or presence (15 mM) of Cyst.

## Supporting information

CyuR RNA Seq

EcoliDEGs

## Acknowledgements

This work was supported by National Institutes of Health (NIH) grants U01AI124316 and GM077402, and Novo Nordisk Foundation Grant Number NNF10CC1016517. We would like to thank Marc Abrams for editing the manuscript.

## Data availability

The RNA-seq raw data files were uploaded to Gene Expression Omnibus (https://www.ncbi.nlm.nih.gov/geo) under an accession id, GSE215167. A token for review is upkhygcedxkvtqf.

## References

1. Durão, P., Balbontín, R. & Gordo, I. Evolutionary Mechanisms Shaping the Maintenance of Antibiotic Resistance. Trends Microbiol. 26, 677–691 (2018).

2. Dwyer, D. J. et al. Antibiotics induce redox-related physiological alterations as part of their lethality. Proc. Natl. Acad. Sci. U. S. A. 111, E2100–9 (2014).

3. Kohanski, M. A., Dwyer, D. J., Hayete, B., Lawrence, C. A. & Collins, J. J. A common mechanism of cellular death induced by bactericidal antibiotics. Cell 130, 797–810 (2007).

4. Shatalin, K., Shatalina, E., Mironov, A. & Nudler, E. H2S: a universal defense against antibiotics in bacteria. Science 334, 986–990 (2011).

5. Shatalin, K. et al. Inhibitors of bacterial H_2_S biogenesis targeting antibiotic resistance and tolerance. Science 372, 1169–1175 (2021).

6. Touati, D. Iron and oxidative stress in bacteria. Arch. Biochem. Biophys. 373, 1–6 (2000).

7. Ono, K. et al. Cysteine Hydropersulfide Inactivates β-Lactam Antibiotics with Formation of Ring-Opened Carbothioic S-Acids in Bacteria. ACS Chem. Biol. 16, 731–739 (2021).

8. Shimada, T., Tanaka, K. & Ishihama, A. Transcription factor DecR (YbaO) controls detoxification of L-cysteine in Escherichia coli. Microbiology 162, 1698–1707 (2016).

9. Zhou, Y. & Imlay, J. A. Escherichia coli Uses a Dedicated Importer and Desulfidase To Ferment Cysteine. MBio 13, e0296521 (2022).

10. Gao, Y. et al. Systematic discovery of uncharacterized transcription factors in Escherichia coli K-12 MG1655. Nucleic Acids Res. 46, 10682–10696 (2018).

11. Li, K. et al. Escherichia coli Uses Separate Enzymes to Produce H2S and Reactive Sulfane Sulfur From L-cysteine. Front. Microbiol. 10, 298 (2019).

12. Loddeke, M. et al. Anaerobic Cysteine Degradation and Potential Metabolic Coordination in Salmonella enterica and Escherichia coli. J. Bacteriol. 199, (2017).

13. Méndez, J. et al. A novel cdsAB operon is involved in the uptake of L-cysteine and participates in the pathogenesis of Yersinia ruckeri. J. Bacteriol. 193, 944–951 (2011).

14. Sastry, A. V. et al. The Escherichia coli transcriptome mostly consists of independently regulated modules. Nat. Commun. 10, 5536 (2019).

15. Lamoureux, C. R., et al. A multi-scale transcriptional regulatory network knowledge base for Escherichia coli. bioRxiv 2021.04.08.439047 (2022) doi:10.1101/2021.04.08.439047.

16. Bethke, J. H. et al. Environmental and genetic determinants of plasmid mobility in pathogenic *Escherichia coli*. Sci Adv 6, eaax3173 (2020).

17. Sastry, A. V. et al. Independent component analysis recovers consistent regulatory signals from disparate datasets. PLoS Comput. Biol. 17, e1008647 (2021).

18. Lim, H. G. et al. Machine-learning from Pseudomonas putida KT2440 transcriptomes reveals its transcriptional regulatory network. Metab. Eng. 72, 297–310 (2022).

19. Ferrández, A. et al. Catabolism of phenylacetic acid in Escherichia coli. Characterization of a new aerobic hybrid pathway. J. Biol. Chem. 273, 25974–25986 (1998).

20. Iwadate, Y. & Kato, J.-I. Identification of a Formate-Dependent Uric Acid Degradation Pathway in Escherichia coli. J. Bacteriol. 201, (2019).

21. Luhachack, L., Rasouly, A., Shamovsky, I. & Nudler, E. Transcription factor YcjW controls the emergency H2S production in E. coli. Nat. Commun. 10, 2868 (2019).

22. Seo, S. W., Kim, D., Szubin, R. & Palsson, B. O. Genome-wide Reconstruction of OxyR and SoxRS Transcriptional Regulatory Networks under Oxidative Stress in Escherichia coli K-12 MG1655. Cell Rep. 12, 1289–1299 (2015).

23. Rodionov, D. A., Gelfand, M. S., Mironov, A. A. & Rakhmaninova, A. B. Comparative approach to analysis of regulation in complete genomes: multidrug resistance systems in gamma-proteobacteria. J. Mol. Microbiol. Biotechnol. 3, 319–324 (2001).

24. Cohen, S. P., Hächler, H. & Levy, S. B. Genetic and functional analysis of the multiple antibiotic resistance (mar) locus in Escherichia coli. J. Bacteriol. 175, 1484–1492 (1993).

25. Rychel, K. et al. iModulonDB: a knowledgebase of microbial transcriptional regulation derived from machine learning. Nucleic Acids Res. 49, D112–D120 (2021).

26. Rodionov, D. A. Comparative genomic reconstruction of transcriptional regulatory networks in bacteria. Chem. Rev. 107, 3467–3497 (2007).

27. Cho, B.-K., Kim, D., Knight, E. M., Zengler, K. & Palsson, B. O. Genome-scale reconstruction of the sigma factor network in Escherichia coli: topology and functional states. BMC Biol. 12, 4 (2014).

28. Rodionova, I. A. et al. A novel bifunctional transcriptional regulator of riboflavin metabolism in Archaea. Nucleic Acids Res. 45, 3785–3799 (2017).

29. Kerr, I. D., Jones, P. M. & George, A. M. Multidrug efflux pumps: the structures of prokaryotic ATP-binding cassette transporter efflux pumps and implications for our understanding of eukaryotic P-glycoproteins and homologues. FEBS J. 277, 550–563 (2010).

30. Matsuo, T. et al. SmdAB, a heterodimeric ABC-Type multidrug efflux pump, in Serratia marcescens. J. Bacteriol. 190, 648–654 (2008).

31. Herman, A. et al. The Bacterial iprA Gene Is Conserved across Enterobacteriaceae, Is Involved in Oxidative Stress Resistance, and Influences Gene Expression in Salmonella enterica Serovar Typhimurium. J. Bacteriol. 198, 2166–2179 (2016).

32. Choe, D. et al. RiboRid: A low cost, advanced, and ultra-efficient method to remove ribosomal RNA for bacterial transcriptomics. PLoS Genet. 17, e1009821 (2021).

33. Lim, H. G. et al. Generation of Pseudomonas putida KT2440 Strains with Efficient Utilization of Xylose and Galactose via Adaptive Laboratory Evolution. ACS Sustainable Chem. Eng. 9, 11512– 11523 (2021).

34. Langmead, B. & Salzberg, S. L. Fast gapped-read alignment with Bowtie 2. Nat. Methods 9, 357–359 (2012).

35. Love, M. I., Huber, W. & Anders, S. Moderated estimation of fold change and dispersion for RNA-seq data with DESeq2. Genome Biol. 15, 550 (2014).

36. Overbeek, R. et al. The SEED and the Rapid Annotation of microbial genomes using Subsystems Technology (RAST). Nucleic Acids Res. 42, D206–14 (2014).

37. Crooks, G. E., Hon, G., Chandonia, J.-M. & Brenner, S. E. WebLogo: a sequence logo generator. Genome Res. 14, 1188–1190 (2004).

38. Mironov, A. A., Vinokurova, N. P. & Gel’fand, M. S. [Software for analyzing bacterial genomes]. Mol. Biol. 34, 253–262 (2000).

39. Kitagawa, M. et al. Complete set of ORF clones of Escherichia coli ASKA library (a complete set of E. coli K-12 ORF archive): unique resources for biological research. DNA Res. 12, 291–299 (2005).

40. Rodionova, I. A. et al. The Nitrogen Regulatory PII Protein (GlnB) and -Acetylglucosamine 6-Phosphate Epimerase (NanE) Allosterically Activate Glucosamine 6-Phosphate Deaminase (NagB) in Escherichia coli. J. Bacteriol. 200, (2018).

41. Rodionova, I. A. et al. Genomic distribution of B-vitamin auxotrophy and uptake transporters in environmental bacteria from the Chloroflexi phylum. Environ. Microbiol. Rep. 7, 204–210 (2015).

42. Sastry, A. V. et al. Machine Learning of Bacterial Transcriptomes Reveals Responses Underlying Differential Antibiotic Susceptibility. mSphere 6, e0044321 (2021).

